# Enhanced expression of Dystrophin, IGF-1, CD44 and MYH3 in plasma and skeletal muscles including Diaphragm of *mdx* mice after oral administration of Neu REFIX Beta 1,3-1,6 glucan

**DOI:** 10.1101/2023.06.06.543858

**Authors:** Senthilkumar Preethy, Shuji Sakamoto, Takuma Higuchi, Koji Ichiyama, Naoki Yamamoto, Nobunao Ikewaki, Masaru Iwasaki, Vidyasagar Devaprasad Dedeepiya, Subramaniam Srinivasan, Kadalraja Raghavan, Mathaiyan Rajmohan, Rajappa Senthilkumar, Samuel JK Abraham

**Affiliations:** Fujio-Eiji Academic Terrain (FEAT), Nichi-In Centre for Regenerative Medicine (NCRM), Chennai, India; Laboratory of Molecular Biology, Science Research Center, Kochi University, Nankoku, Japan; Antony- Xavier Interdisciplinary Scholastics (AXIS), GN Corporation Co. Ltd., Kofu, Japan; Genome Medical Sciences Project, National Center for Global Health and Medicine, Konodai, Ichikawa, Chiba, Japan; Dept. of Medical Life Science, Kyushu University of Health and Welfare, Nobeoka, Japan; Institute of Immunology, Junsei Educational Institute, Nobeoka, Miyazaki, Japan; Centre for Advancing Clinical Research (CACR), University of Yamanashi - School of Medicine, Chuo, Japan; Mary-Yoshio Translational Hexagon (MYTH), Nichi-In Centre for Regenerative Medicine (NCRM), Chennai, India; Dept of Paediatric Neurology, Jesuit Antonyraj memorial Inter-disciplinary Centre for Advanced Recovery and Education (JAICARE), Madurai, India; R & D, Sophy Inc., Japan; Levy-Jurgen Transdisciplinary Exploratory (LJTE), Global Niche Corp., Wilmington, DE, USA; Haraguchi-Parikumar Advanced Remedies (HARP), SoulSynergy Ltd., Phoenix, Mauritius

**Keywords:** β-glucan, Duchenne Muscular Dystrophy (DMD), *mdx* mice, CD44, MYH3

## Abstract

**Introduction:** Duchenne muscular dystrophy (DMD) is a rare genetic disease, causing muscle degeneration due to lack of dystrophin with inadequate muscle regeneration culminating in muscle dysfunction. The N-163 strain of *Aureobasidium Pullulans* produced Beta-1,3-1,6-glucan (Neu REFIX) reported to be safe with anti-inflammatory and anti-fibrotic efficacy earlier, herein we evaluated its effects on muscle regeneration in *mdx* mice.

**Methods:** Forty-five mice in three groups (n=15 each): Group 1 (normal), Group 2 (*mdx* control), and Group 3 (*mdx* fed Neu REFIX) were evaluated for 45 days. IGF-1, Dystrophin, CD44 and MYH3 in diaphragm, plasma and skeletal muscle were evaluated by ELISA and immunohistochemistry.

**Results:** Mean IGF-1 expression was 20.32% and 16.27% higher in plasma (p = 0.03) and diaphragm respectively in Neu-REFIX group. Mean dystrophin was higher in Neu-REFIX group by 70.3% and 4.7% in diaphragm and plasma respectively than control. H-score intensity of CD44+ was >2.0 with an MYH3-positivity 20% higher in Neu-REFIX than control.

**Conclusion:** Oral administration of Neu REFIX was safe. Significantly enhanced plasma IGF-1 beside increased Dystrophin, MYH3 and CD44, proving a restoration of muscle regeneration and differentiation, especially in diaphragm, makes us recommend it as a disease modifying adjuvant in both early and advanced stages of DMD.

## INTRODUCTION

Dystrophin, a protein that gives the membrane of muscle fibres structural rigidity and integrity, when encoded by a faulty gene leads to Duchenne muscular dystrophy (DMD), an X-linked hereditary severe form of muscular dystrophy [1]. The lack of dystrophin in muscle fibres causes membrane fragility, a high risk of injury, severe inflammation, and oxidative stress, leading to muscle necrosis. Continuous damage result in fibrosis of the muscles and their dysfunction leading to loss of ambulation followed by pre-mature death in the early 20s or 30s due to cardiac and respiratory failure [1]. In muscular dystrophy, though the satellite stem cells are activated to regenerate in response to injury, the invasion of immune cells, primarily macrophages occur together with inflammation leading to improper muscle maturation and progressive replacement by fibrous-fatty tissue ensues [2].

Although a lack of dystrophin is the primary cause of the manifestations of DMD, ensuing chronic inflammation seems to be the key factor curtailing proper muscle regeneration [1]. While there is no definitive cure for the disease, approaches to control inflammation by use of corticosteroids has been a standard of care. However, due to the associated adverse effects of corticosteroids, their continuous administration in all the patients is quite difficult. While gene therapies are in clinical trials, even the approved exon skipping therapies with some benefits in basic and translational studies, haven’t yet yielded a clinically satisfactory outcome, which may be attributed to insufficient amelioration of inflammation, fibrosis, apart from limited muscle regeneration and maturation [3].

In this scenario, there is need for a safe and easily administrable disease modifying approach. We have earlier demonstrated the anti-inflammatory, anti-fibrotic and immune-modulating effects of a biological response modifier B-glucan (BRMG) produced by the N-163 strain of *Aureobasidium pullulans* (*A.pullulans*) in pre-clinical *mdx* mice model [4] and human clinical studies of DMD [5]. In the *mdx* animal study done earlier, this B-Glucan was able to ameliorate inflammation proven by a significant decrease in inflammation score and fibrosis, levels of plasma ALT, AST, LDH, IL-13, and haptoglobin, and an increase in levels of anti-inflammatory TGF-β [4]. In the human clinical study done in 27 patients, anti-inflammatory effects proven by a decrease in IL-6 levels, and anti-fibrotic effects by a decrease in TGF-β levels apart from improved muscle strength and increased plasma dystrophin levels [5] were demonstrated. However, with an objective to evaluate the effects of this N-163 B-Glucan food supplement on muscle regeneration and differentiation, this study was undertaken in *mdx* mice model of DMD.

## METHODS

The guidelines for reporting in vivo animal experiments were followed when conducting this investigation. Male, three-week-old C57BL/10SnSlc mice were purchased from Japan SLC, Inc. (Japan). Male, three-week-old C57BL/10-*mdx*/Jcl mice were purchased from CLEA Japan (Japan). The animals were treated with care in accordance with the Ministry of the Environment’s Act No. 105 of 1 October 1973’s Act on Welfare and Management of Animals, its Notice No. 88 of April 28, 2006’s Standards Relating to the Care and Management of Laboratory Animals and Relief of Pain, and its Guidelines for Appropriate Conduct of Animal Experiments.

Forty-five mice were included in the study [4] and based on their body weight, the day before the start of treatment were randomized into the following three groups of 15 mice each. Randomisation was performed by body weight-stratified random sampling using Excel software.

Group 1: Normal- Fifteen C57BL/10SnSlc mice were without any treatment until sacrifice.

Group 2: Vehicle- Fifteen *mdx* mice were orally administered vehicle [pure water] at a volume of 10 mL/kg once daily from days 0 to 45.

Group 3: N-163 β-glucan- Fifteen *mdx* mice were orally administered with N-163 strain produced B-glucan (Neu-REFIX^TM^), was provided by GN Corporation Co. Ltd at a dose of 3 mg/kg as API in a volume of 10 mL/kg once daily for 45 days.

The animals were housed in TPX cages (CLEA Japan) and maintained in an SPF facility under controlled conditions of temperature, humidity, lighting (12-hour artificial light and dark cycles and air exchange. Viability, clinical signs (lethargy, twitching, and laboured breathing), and behaviour were monitored daily. Body weight was recorded daily before treatment. The mice were observed for significant clinical signs of toxicity, morbidity, and mortality before and after administration. The animals were sacrificed on day 45 by exsanguination through the abdominal vena cava under isoflurane anaesthesia (Pfizer Inc.). After sacrifice, the quadriceps, gastrocnemius, soleus, plantaris, tibialis anterior, extensor digitorum longus, diaphragm, and myocardium muscles were collected. Individual muscle weights were measured. Each muscle was separated, dissected, and stored for analysis [4].

The expression of IGF-1 in diaphragm muscle and plasma from (n=8 mice in each group) mice was analyzed using an ELISA kit (Mouse/Rat IGF1 Sandwich ELISA Kit-Proteintech, Catalogue Number: KE10032). All reagents were brought to room temperature before use, and care was taken to avoid cross-contamination by changing pipette tips between each addition of standard, sample, and reagent, as well as using separate reservoirs for each reagent. The required number of microplate strips was taken out and excess strips were returned to the foil pouch with a drying reagent pack, resealed, and stored at 4°C for use within one week. The microplate layout was preset to include control, standard, and sample groups, with 100 μL of each standard and sample added to the appropriate wells, ensuring uninterrupted addition completed within 5 to 10 minutes. All standards, controls, and samples were assayed in duplicate. The plate was sealed with a cover seal and incubated for 2 hours at 37°C. After incubation, the cover seal was gently removed, and the liquid was discarded by aspirating or decanting. Residual solution was removed by tapping the plate on fresh paper towels. The wells were washed four times with 1X Wash Buffer, using 350-400 μL per well, and the plates were tapped firmly on fresh towels 10 times to remove residual Wash Buffer. 100 μL of 1X Detection Antibody solution was added to each well, the plate was sealed with a cover seal, and incubated for 1 hour at 37°C. The wash step was repeated as previously described. 100 μL of 1X Streptavidin-HRP solution was added to each well, the plate was sealed with a cover seal, and incubated for 40 minutes at 37°C. The wash step was repeated again. For signal development, 100 μL of TMB substrate solution was added to each well, protected from light, and incubated for 15 to 20 minutes until the substrate solution remained colourless. Colour development was quenched by adding 100 μL of Stop Solution to each well in the same order as the TMB substrate addition, mixing gently by tapping the side of the plate. Absorbance was read immediately after adding the Stop Solution on a microplate reader at 450 nm. Data analysis involved calculating the average of duplicate readings (OD values) for each standard and sample, subtracting the average of the zero standard absorbance (delta OD values). The concentration versus δ absorbance (delta OD values) was calculated from the calibration curve.

Dystrophin expression in the diaphragm (n=8 mice in each group) was quantified using the SimpleStep ELISA® (Enzyme-Linked Immunosorbent Assay) kit, designed for the measurement of dystrophin protein in human and mouse cell and tissue extract samples. The procedure was performed according to the manufacturer’s instructions. Diaphragm muscle extracts were prepared as per standard protocols. Briefly, samples were homogenized in lysis buffer, centrifuged to remove debris, and the supernatants were collected for analysis. For the ELISA Procedure, diluted samples and standards were prepared in the provided diluent according to the recommended concentrations. 50 µL of standards or samples, a cocktail of capture and detection antibodies (enzyme-conjugated antibodies) were added to each well of an ELISA plate precoated with the capture antibody.

The plate was incubated for 1 hour at room temperature on a plate shaker set to 400 rpm. After incubation, wells were washed three times with 350 µL of 1x wash buffer to remove unbound materials. 100 µL of TMB Development Solution was added to each well. The plate was then incubated for 10 minutes in the dark at room temperature. The reaction was stopped by adding 100 µL of Stop Solution to each well. The absorbance was measured at 450 nm using a microplate reader. The absorbance values at 450 nm were used to construct a standard curve from the standards provided. Dystrophin concentrations in the samples were calculated from this standard curve. All samples and standards were assayed in duplicate to ensure accuracy and reproducibility.

For dystrophin expression analysis in plasma, mouse plasma samples (n=8 mice in each group) (25x dilution) were fixed in 96-well microplates and incubated overnight at 4°C. After fixation, a rabbit anti-dystrophin polyclonal antibody (400x dilution) was added to each well and incubated for 1 hour. This was followed by the addition of a biotin-labelled goat anti-rabbit antibody (20,000x dilution) and POD-labelled avidin (10,000x dilution). For signal development, 100 μL of TMB substrate solution was added to each well, and incubated for 15 to 20 minutes. Colour development was quenched by adding 100 μL of Stop Solution to each well. Absorbance was read immediately after adding the Stop Solution on a microplate reader at 450 nm. Data analysis involved calculating the average of duplicate readings (OD values) for each standard and sample, subtracting the average of the zero standard absorbance (delta OD values). The results were expressed as δOD values. The ELISA was performed using a simple, direct method to detect dystrophin levels.

For CD44 (n=6 mice in each group) and MYH3 (n=9 mice in each group) expression, muscle preparation for histological analysis was done by placing the tragacanth gum on cork disk, with only enough tragacanth gum to provide foundation for the oriented muscle. The tragacanth gum was placed on one end so that the long axis of the muscle is perpendicular to the cork disc. The specimen was rapidly frozen and placed into isopentane cooled in liquid nitrogen. The frozen block was transferred on to dry ice and the isopentane was evaporated for around one hour and stored at −80 °Cross sections were cut from paraffin blocks of the muscle (TBD) tissue using a rotary microtome (Leica Microsystems). After sectioning, each slide was coded as a number for blind evaluation. Each number was generated using the RAND function of the Excel software, sorted in ascending order, and assigned to the slides. The tissue slides were used for staining. For immunohistochemistry staining of CD44, sections were cut from the frozen blocks and fixed in acetone. Endogenous peroxidase activity was blocked using 0.03% H2O2 for 5 minutes followed by incubation with Block Ace (DaiNippon Sumitomo Pharma Co.Ltd., Japan) for 10 minutes. The sections were incubated with antibody against CD44 (Rabbit Monoclonal Anti-CD44(E7K2y) purchased from Cell Signalling Technology (Danvers, MA, USA). For immunofluorescence staining of Myosin heavy chain (MYH3), the frozen sections were fixed in freshly prepared 4% paraformaldehyde for 15 min at room temperature and were then permeabilized with 1% Triton X-100 in PBS for 20 min at room temperature. The sections were blocked with blocking solution (0.2% gelatin, 1% BSA and 0.05% Tween 20 in PBS) for 30 min at room temperature. After blocking, the sections were immunostained with primary antibodies diluted in blocking solution at 4 °C overnight. After washing with TBS containing 0.05% Tween 20 (TBST), the sections were incubated with secondary antibodies diluted in blocking solution for 2h at room temperature. After washing with TBST, the sections were stained with DAPI to label nuclei. The stained sections were mounted with ProLong Gold antifade reagent (Invitrogen, USA). The following antibody was used: Myh3 (sc-53091, Santa Cruz Biotechnology, 1:100). Images were taken with all-in-one fluorescence microscope BZ-X800 (Keyence, Japan). The positive areas of those biomarkers were evaluated using H-score [6] & SketchAndCalc™ software, respectively.

Statistical analyses were performed using the Prism Software 6 (GraphPad Software, USA). Statistical analyses were performed using ANOVA and Unpaired T tests. Comparisons were made between the following groups: 1) Group 2 (Vehicle) vs. Group 1 (Normal) and Group 3 (N-163 β-glucan). Statistical significance was set at P < 0.05. Results are expressed as mean ± SD.

## RESULTS

The mean body weight of mice in the vehicle group was significantly lower than that of mice in the normal group but there was no significant difference in the mean body weight between the vehicle group and the N-163 β-glucan.

In plasma, IGF-1 expression was 4145 ± 620.4 pg/ml in Gr.1 (WT), which decreased to 3366 ± 645.8 pg/ml in Gr.2 (*mdx*-control). Gr.3 (*mdx-*Neu REFIX) displayed a statistically significant 20.32% increase in plasma IGF-1 compared to Gr.2 (*mdx-*control), with a mean level of 4050 ± 602.2 pg/ml (p = 0.03) (Figure 1). Mean level of IGF-1 expression in diaphragm muscle in Gr.1(WT) was 173.6 ± 74.5 pg/ml, while Gr.2 (*mdx*-control) exhibited a decrease to 107.9 ± 25.11 pg/ml. In Gr.3 (*mdx*-Neu REFIX), IGF-1 levels were 125.4 ± 23.47 pg/ml, the difference between the three groups was statistically significant (p=0.04). The levels of IGF-1 in diaphragm was 16.27% higher in Gr.3 (*mdx*-Neu REFIX) than Gr.2 (*mdx*-control), though the difference was not statistically significant (p = 0.13).

**Figure 1:**
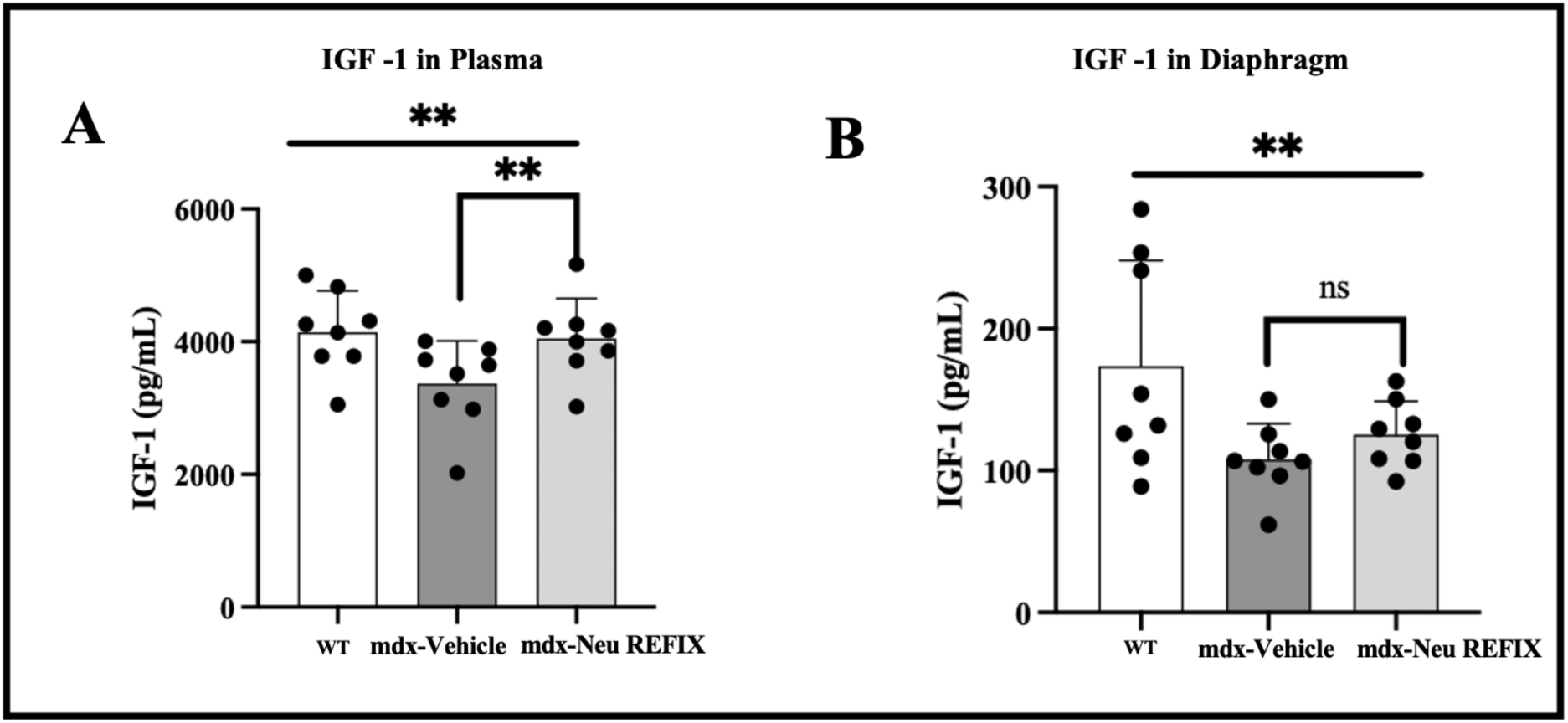
IGF-1 expression in A. Plasma and B. Diaphragm showing increased expression in Neu REFIX compared to the Vehicle group.

The mean level of dystrophin expression in plasma was (OD value: 0.6223 ± 0.0515) in Gr.1 (WT), decreasing to (OD value: 0.5504 ± 0.0319 in Gr.2 (*mdx*-control). In Gr.3 (*mdx*-Neu REFIX), plasma dystrophin was (OD value: 0.5763 ± 0.0490) (ANOVA p=0.01), 4.7% higher than Gr.2 (*mdx*-control), but this increase was also not statistically significant (p = 0.2). The mean dystrophin expression in diaphragm muscle was 201.7 ± 126.5 pg/ml in Gr.1 (WT), which decreased to 87.65 ± 58.4 pg/ml in Gr.2 (*mdx*-control). In Gr.3 (*mdx*-Neu REFIX), the expression was 149.3 ± 92 pg/ml, representing a 70.3% increase compared to Gr.2, though this difference was not statistically significant (p = 0.08) compared to Gr.2 (Figure 2).

**Figure 2:**
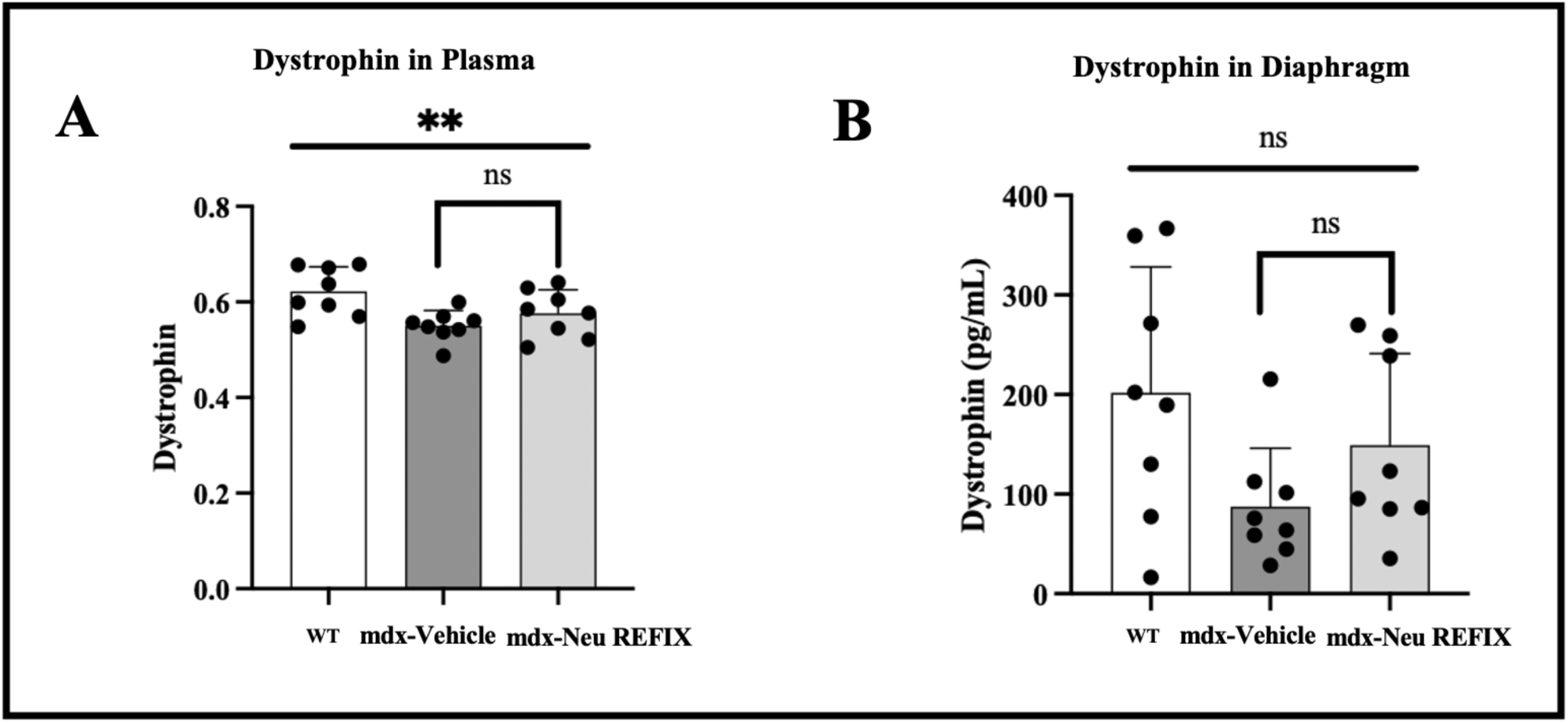
Dystrophin expression in A. Plasma and B. Diaphragm showing increased expression in Neu REFIX compared to the Vehicle group.

The H-score is one of the commonly employed endpoints to gauge biomarker expression. Individual cells are first identified using the H-score algorithm and then the cells are categorised as either positive or negative depending on the relative expression of the biomarker of interest. Based on the strength of the biomarker signal, the positive cells are further divided into high (3+), medium (2+), or low (1+) categories [6]. The proportion of the weighted sum of the number of positive cells to the total number of detected cells determines the H-score. The H-score extracts the biomarker of interest’s intensity and percentage from the IHC image. The H-score for CD44 expression was 0.1 ± 0.3 in Gr. 1 (Normal), 1 ± 0.18 in Gr. 2 (Vehicle) and 2.16 ± 0.89 in Gr. 3 (N-163), thus a significantly higher H-score in the Neu-REFIX group compared to Gr.2 and Gr.1 (p-value = 0.0002) (Figure 3). SketchAndCalc™ is an irregular area calculator app for all manner of images containing irregular shapes. When calculated using SketchAndCalc, the MYH3 positive area was 1.72 cm^2^ with a total area of positively stained cells being 256 units and percentage of positive region was 0.665 in Gr.1. In Gr. 2, the MYH3 positive area was 10.78 cm^2^ with a total area of positively stained cells being 272.6 units and percentage of positive region was 3.954. In Gr. 3 (Neu-REFIX) the MYH3 positive area was highest among the groups, 15.11 cm^2^ with a total area of positively stained cells being 305.25 units and percentage of positive region was 4.954 (Figure 4)

**Figure 3:**
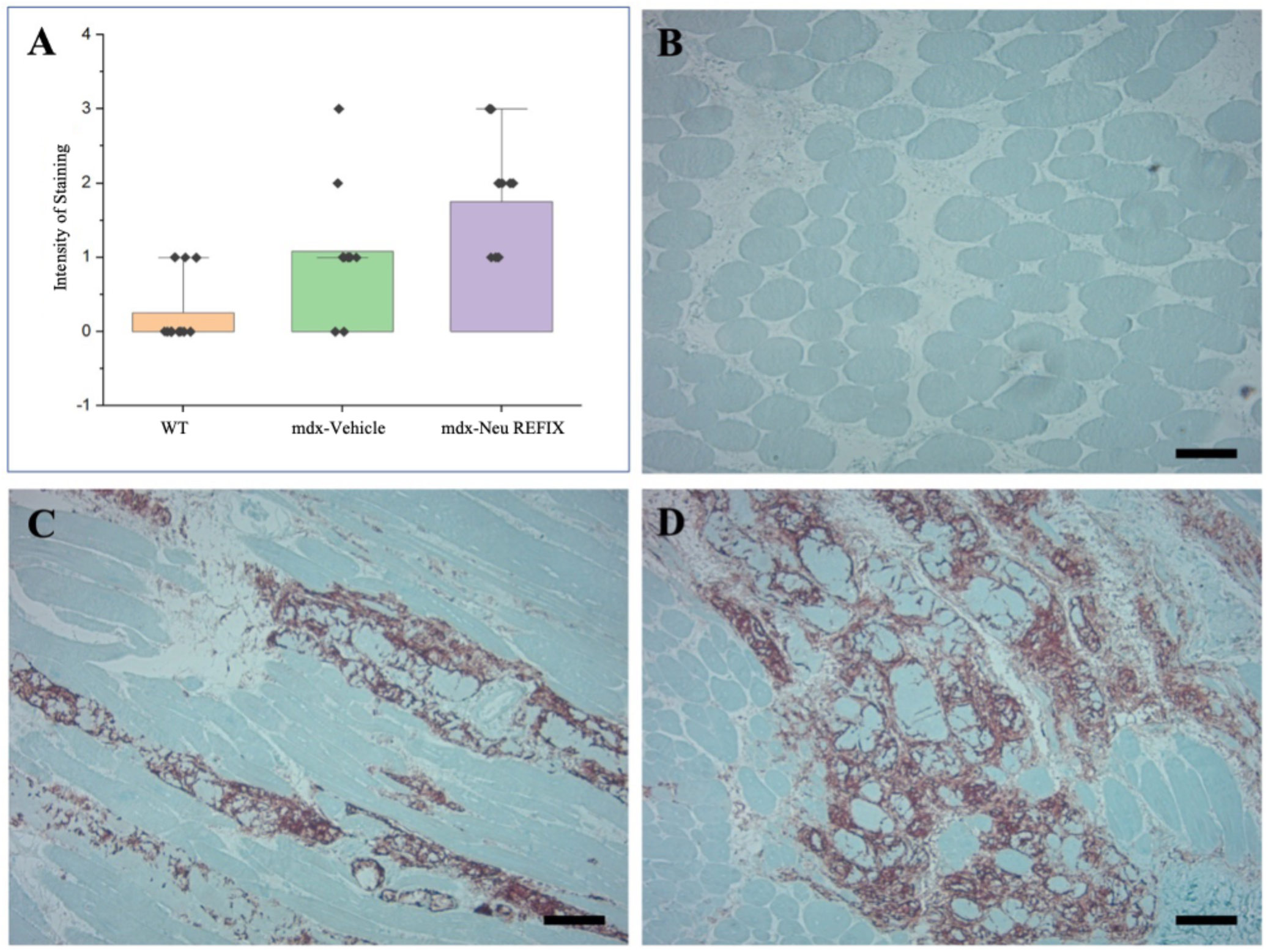
Immunohistochemistry staining of CD44 in muscle of *mdx* mice; A. Comparison of H-SCORE between the groups; B. (Gr. 1) Normal mice; C. (Gr.2) *mdx* mice - Vehicle group and D. (Gr.3) *mdx* mice –Neu REFIX group. Scale bar: 100 mm

**Figure 4:**
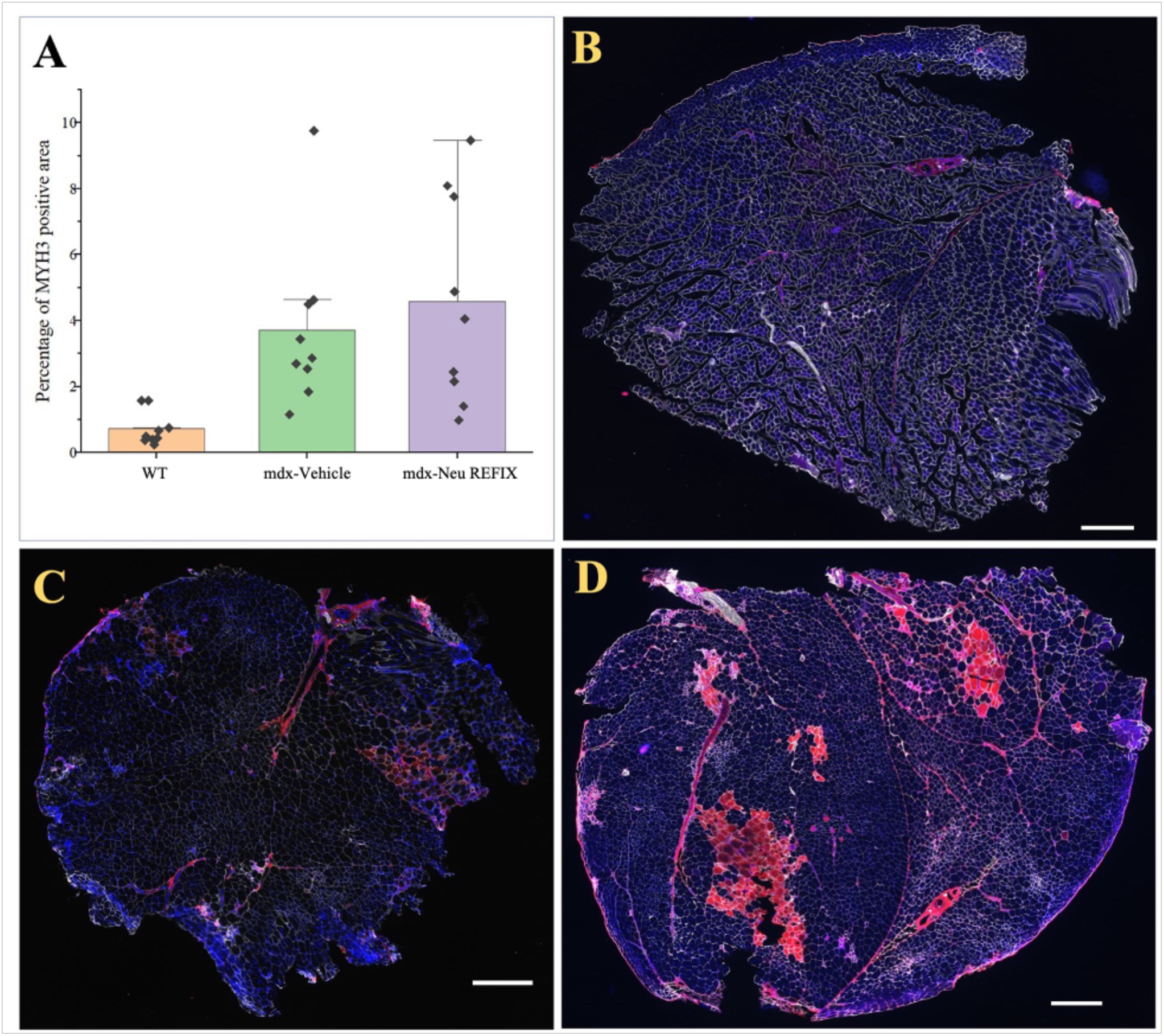
A. Percentage of MYH3 positive fibres in the three groups; B. (Gr. 1) Normal mice; C. (Gr.2) *mdx* mice - Vehicle group and D. (Gr.3) *mdx* mice – Neu REFIX group; Scale bar: 100 mm

## DISCUSSION

Duchenne Muscular Dystrophy (DMD) remains a challenging disorder with limited therapeutic options. While gene therapy and exon-skipping treatments have shown promise, they often come with adverse effects and are not applicable to all DMD patients. Given this gap, we explored a novel disease-modifying approach using Neu REFIX, a β-glucan-based biological response modifier. Previous pre-clinical studies in Sprague Dawley rats [7] and non-alcoholic steatohepatitis (NASH) animal models [8], as well as clinical trials involving healthy volunteers [9] and COVID-19 [10] patients, have demonstrated the anti-inflammatory and anti-fibrotic properties of Neu REFIX. In a clinical study involving DMD patients [5], we observed a decrease in inflammatory markers such as IL-6 and TGF-β, alongside functional improvements in muscle strength and plasma dystrophin levels. In a follow-up study in *mdx* mice [4], we found a reduction in inflammation scores, fibrosis, and serum levels of ALT, AST, and LDH, as well as increases in anti-inflammatory TGF-β levels and balanced regulation of centrally nucleated fibres. Having established its anti-inflammatory properties, we then sought to examine the effects of N-163 β-glucan on muscle regeneration and maturation for the same duration (45 days) in the present study.

Current therapies for DMD, such as stem cell injections, exon-skipping therapies, and gene therapies, may have limited efficacy in an inflammatory environment. Moreover, gene therapy itself may trigger immune-mediated inflammation [11], potentially exacerbating the disease. To be considered effective, a DMD therapy must address two critical aspects: (i) reduction of inflammation and fibrosis associated with dystrophin deficiency and (ii) promotion of muscle stem cell regeneration and maturation into functional muscle cells. Although muscle injury can activate tissue-resident satellite stem cells, myofiber formation and maturation are often hindered by chronic inflammation. Therefore, reducing inflammation is crucial for any regenerative approach in DMD. Our approach with Neu REFIX offers a multi-modal benefit for DMD. In the current study, Neu REFIX demonstrated an increase in dystrophin levels in plasma and diaphragm muscle, along with elevated levels IGF-1 and increased markers of muscle regeneration (CD44 and MYH3). Additionally, the agent has a favourable safety profile established in clinical studies [5, 9,10], making it potentially suitable for a broad age range and disease stages in DMD patients.

In our prior 45-day clinical trial with DMD patients [5], we observed a restoration of plasma dystrophin expression; however, muscle-specific expression could not be studied due to challenges in obtaining repeated muscle biopsies. In the current study, Neu REFIX-treated *mdx* mice (Gr.3) exhibited higher dystrophin levels in muscle tissue compared to the *mdx*-control group (Gr.2), In earlier trials with definitive therapies, such as gene therapy and exon-skipping approaches, dystrophin expression ranged from 20% to 92% over a relatively short duration of 10–30 days [12–14]. However, it should be noted that, in our study, Neu REFIX is an adjunct disease modifying treatment has resulted in a dystrophin expression nearly 70% higher than that observed in *mdx*-vehicle or *mdx*-control mice, providing proof of concept for its potential as an adjunct therapy in DMD. Additionally, while previous exon-skipping studies have shown that dystrophin expression alone may not always result in functional improvement due to partial functionality [13, 15], our previous human trial demonstrated improved muscle function with Neu REFIX [16], suggesting that this β-glucan formulation may enhance dystrophin increase that correlates with improved muscle functionality.

The therapeutic effects of Neu REFIX in the present study may be attributed to its potent anti-inflammatory activity, which has been shown to create a favorable tissue microenvironment for muscle regeneration. Inflammatory pathways are critical in the progression of DMD, as chronic inflammation exacerbates fibrosis and impedes the restoration of dystrophin [17]. By mitigating inflammation, Neu REFIX may facilitate the resolution of fibrosis [8] and enhance the muscle’s regenerative capacity. Previous studies in the NASH animal model have also demonstrated that Neu REFIX exerts its effects through epigenetic modulation mediated by the gut microbiome [18]. The beneficial effects on gut microbiome reconstitution was observed in the clinical study in DMD as well [19]. This mechanism, coupled with its proven safety profile, suggests a broader potential for Neu REFIX in addressing the complex pathology of DMD. However, further mechanistic studies are essential to delineate these pathways and confirm the role of Neu REFIX in promoting dystrophin restoration and muscle repair.

IGF-1 is essential for muscle repair and regeneration as it promotes the proliferation and differentiation of skeletal muscle cells. Recombinant IGF-1 therapy has shown limited effects in enhancing motor function in DMD patients [20], and transgenic overexpression in *mdx* mice increased fibre size [21] but did not alter muscle IGF-1 concentration. In contrast, Neu REFIX administration led to increased IGF-1 levels in both muscle and plasma in the present study. The natural stimulation of IGF-1 through Neu REFIX without external administration indicates its potential for enhancing muscle regeneration and function in DMD. The evaluation of IGF-1 levels in the serum provides significant diagnostic value in assessing muscle regeneration and differentiation. However, a comprehensive assessment of treatment efficacy necessitates the inclusion of additional critical indicators, such as serum creatine kinase (CK) levels and functional assessments like muscle force which we are planning to evaluate in our future studies.

MYH3 and CD44 expression levels are indicative of muscle regeneration and maturation. MYH3 is a neonatal myosin expressed during early muscle development and re-expressed during muscle repair [22]. In our study, a 20% increase in MYH3 expression in Neu REFIX-treated mice validated enhanced muscle regeneration and stem cell activation. Additionally, CD44 expression, a marker associated with myogenic cell differentiation [23], was significantly higher in the Neu REFIX group, indicating proper differentiation and maturation of muscle fibres.

The gut-muscle axis plays a critical role in inflammation regulation, and improvements in gut microbiome composition have been linked to reduced inflammation [24]. In our previous study on DMD patients treated with N-163 β-glucan (Neu REFIX), we observed significant beneficial microbiome reconstitution [19] associated with reduced inflammatory markers. This connection between gut health and muscle recovery is supported by studies showing that a balanced gut microbiome can modulate systemic inflammation, which is relevant for the anti-inflammatory and anti-fibrotic effects seen in our study.

While this study provides promising evidence for Neu REFIX’s role in muscle regeneration and anti-inflammatory effects in *mdx* mice, validation in human muscle biopsies is required. However, such studies are invasive and challenging to conduct in clinical settings. In our previous human study, improvements in muscle strength and function, as measured by the Medical Research Council (MRC) scale [5, 16], align with the results in this animal model, further supporting the β-glucan’s potential for enhancing muscle regeneration. Future studies should explore the molecular mechanisms behind the observed therapeutic effects, particularly the interactions between dystrophin, IGF-1, and the gut-muscle axis. Additionally, we recommend extending the study duration beyond 45 days to examine the long-term effects of Neu REFIX on dystrophin expression and muscle regeneration. Studies in animal models with satellite cell depletion or aged mice with reduced satellite cell reservoirs would also provide valuable insights into the β-glucan’s impact on muscle stem cell dynamics in DMD.

On the point that different muscle tissues were used for evaluation of the markers in the current study, the rationale is to focus on evaluating the therapeutic potential of Neu REFIX in the diaphragm muscle, given its critical involvement in the advanced stages of DMD [25]. DMD being a progressive disease with several research studies primarily focussing on skeletal muscles, we prioritized the evaluation of the diaphragm muscle in this extended study building upon previous analyses [4] conducted on the same group of animals on different muscle tissues. The diaphragm was chosen due to its clinical relevance in late-stage DMD [25], particularly for patients who may not benefit from exon-skipping or gene therapy, wherein Neu REFIX’s beneficial effects on the diaphragm could be used to better understand its applicability in mitigating disease progression and improving respiratory muscle function in advanced stages of DMD.

This study while encouraging has limitations. One of the limitations is that the study duration was only 45 days which was done to match the duration of the clinical study done earlier [5]. This study primarily aimed to evaluate the safety and preliminary efficacy of oral administration of Neu REFIX β-glucan in mdx mice over a 45-day period. While a significant increase in IGF-1 levels was observed in plasma, other biomarkers did not achieve statistical significance within the study duration. These findings highlight the challenges of demonstrating robust therapeutic effects in a short-term study for a chronic condition like DMD. The limited study duration may have influenced the statistical outcomes, as chronic diseases often require longer intervention periods to manifest significant biological changes. To address this, a six-month clinical study has been earlier conducted [16], and future long-term preclinical studies in mdx mice and clinical trials are being planned. With regard to the *mdx* mice model, it may not exactly recapitulate the human pathology of DMD. Similar studies in severe muscle degenerative models of DMD such as DBA/2-*mdx* [26], DMD-edited micro minipigs [27] etc., need to be undertaken, especially when we propose Neu REFIX as a disease modifying adjuvant for advanced stages of the disease, because these animal models also simulate the cardiac degeneration and other manifestations of DMD in humans. The pro-regenerative effects of Neu REFIX were evidenced by the observed increase in MYH3 expression via immunofluorescence and differentiation markers like CD44. However, the potential risk of excessive regeneration depleting the satellite cell pool warrants careful consideration, as this could impair long-term muscle regenerative capacity. In our previous study [4] we quantified centronucleated fibers to evaluate muscle regeneration after Neu REFIX administration. Building on these findings, we are currently conducting additional functional studies in the *mdx* mouse model. Future analyses will include the evaluation of genes involved in the myogenic cascade and late-stage differentiation markers via RT-PCR. These assessments will provide critical insights into the balance between regeneration and satellite cell pool preservation, further clarifying the long-term therapeutic potential of Neu REFIX. While the observed increase in dystrophin levels in the diaphragm is promising, a limitation of this study is the lack of immunofluorescence analysis to confirm and visualize the spatial localization of the protein. Immunofluorescence provides critical insights into dystrophin distribution within muscle fibers and its functional relevance. Future studies will address this limitation by including immunofluorescence analysis to enhance our understanding of dystrophin expression patterns and their role in muscle function restoration. Another limitation of the current study is the absence of pharmacokinetic analyses to assess the biodistribution of Neu REFIX and its extent of penetration into muscle tissues. Understanding the drug’s pharmacokinetics is essential to correlate its systemic administration with observed therapeutic outcomes. We aim to address this limitation in future studies by conducting detailed pharmacokinetic and biodistribution evaluations to provide a more comprehensive understanding of Neu REFIX’s therapeutic mechanism. After such relevant validations, application of this adjuvant approach may have to be tried in combination with primary treatments such as gene therapy, exon skipping therapy etc. as a disease modifying agent.

## CONCLUSION

Neu REFIX an orally administrable, safe, allergen-free β-1,3-1,6 glucan, has demonstrated its potential as a disease-modifying therapeutic adjuvant for DMD in *mdx* mice model, by enhancing dystrophin and IGF-1 expression along with muscle regeneration markers MYH3 and CD44 in a short duration of 45 days. Having earlier proven for its efficacy as an anti-inflammatory and anti-fibrotic agent and having demonstrated enhanced dystrophin in plasma in a clinical study, we recommend further long term multi-centric clinical studies in patients with DMD. Neu REFIX by its anti-inflammatory potential could become an adjuvant to even exon-skipping and gene therapies, in mitigating their adverse effects and improve therapeutic outcomes along with its standalone efficacy per se adding to such definitive approaches.

## Declarations

### Ethics approval and consent to participate

**All experimental protocols were approved by SMC Laboratories, Japan’s IACUC (Study Protocol no: SP_SLMA143-2208-3). All methods are reported in accordance with ARRIVE guidelines**. This study was conducted in accordance with the Animal Research: Reporting of In Vivo Experiments Guidelines. C57BL/10SnSlc mice (3 weeks of age, male) were obtained from Japan SLC, Inc. (Japan). C57BL/10-*mdx*/Jcl mice (3 weeks of age, male) were obtained from CLEA Japan (Japan). All animals used in this study were cared for following guidelines: Act on Welfare and Management of Animals (Ministry of the Environment, Act No. 105 of October 1, 1973), Standards Relating to the Care and Management of Laboratory Animals and Relief of Pain (Notice No.88 of the Ministry of the Environment, April 28, 2006) and Guidelines for Proper Conduct of Animal Experiments (Science Council of Japan, June 1, 2006).

### Availability of data and material

All data generated or analysed during this study are included in the article itself.

### Author Contribution Statement

S.J.A. and N.I contributed to conception and design of the study. K.I, S.S and T.H helped with the experiments and analysis. R.S and M.R performed the literature search. S.J.A, and S.P. drafted the manuscript. N.Y, M.I, V.D, K.R and S.S performed critical revision of the manuscript. All the authors read and approved the submitted version.

## Acknowledgments

The authors would like to dedicate this paper to the memory of Mr. Takashi Onaka, who passed away on the 1st of June, 2022 at the age of 90 years, who played an instrumental role in successfully culturing and industrial scale up of AFO-202 and N-163 strains of *Aureobasidium pullulans* after their isolation by Prof. Noboru Fujii, producing the novel beta glucans described in this study. They thank, Mr. Yasushi Onaka and Mr. Masato Onaka, of Sophy Inc. for technical clarifications, Ms. Eiko Amemiya of II Dept. of Surgery, University of Yamanashi for secretarial assistance, Dr. Yoshitsugu Aoki and Dr. Katsura Minegishi of Department of Molecular Therapy, National Centre for Neurology and Psychiatry (NCNP), Tokyo, Japan, for their guidance in designing the experiment and training of the animal study experts, Ms. Mami Fujimoto, administrative staff of NCNP for facilitating the collaborative arrangement with GN Corporation Co Ltd., and Ms. Yoko Okubo of NCNP for technical guidance of the histological sample preparations; Ms. Yoshiko Amikura of GN Corporation, Japan for liaising between the institutes and Ms. Kaori Ikewaki for her technical assistance with immunohistochemistry evaluation.

